# Identifying mood instability and circadian rest-activity patterns using digital remote monitoring and actigraphy in participants at risk for bipolar disorder

**DOI:** 10.1101/2025.01.20.633946

**Authors:** Priyanka Panchal, Natalie Nelissen, Niall M. McGowan, Lauren Z. Atkinson, Kate E. A. Saunders, Paul J. Harrison, Matthew Rushworth, Dejan Draschkow, John Geddes, Anna C. Nobre, Catherine J. Harmer

## Abstract

Mood instability and circadian rhythm disruptions are both of increasing interest with regard to a number of psychiatric disorders, notably bipolar disorder (BD), but understanding of their nature and their interrelationship are incomplete. By definition, both have an integral temporal component and, as such, measuring them longitudinally and remotely is desirable.

We conducted the Cognition and Mood Evolution across Time (COMET) study to assess the feasibility and value of digital devices to capture mood, its instability, and daily rest-activity patterns, over a 10-week period, in two groups of participants. The first group (n=37) were selected as scoring ≥7 on the Mood Disorder Questionnaire (MDQ) (‘high MDQ’), thereby having a history of mood elevation and being at risk for BD. They were compared with a group (n=37) scoring ≤5 on the MDQ (‘low MDQ’). Over a 10-week period, using a tablet, mood was rated daily, clinical ratings of depression, mania, and anxiety were captured weekly via the True Colours app, and a GENEactiv actigraph was worn to capture rest-activity pattern data.

The main findings are that (1) MDQ score predicts mood instability; (2) high MDQ score is associated with more negative affect and mood symptoms than people with low MDQ, and with a different circadian activity profile; and (3) mood instability and circadian indices appear uncorrelated.

The implications are that (1) remote monitoring of these domains is feasible and valuable; (2) selection of participants based on MDQ score is useful for studying mood (in)stability; and (3) the approach has potential for studies of clinical populations and for experimental medicine studies assessing interventions to reduce mood instability.

## Introduction

Mood instability, defined as rapid oscillation of intense affect, with difficulty in regulating these oscillations for their behavioural consequences, is a common feature across psychiatric disorders including bipolar disorder. Disruption in circadian rest-activity patterns is often seen alongside these mood changes, but it is not known whether these observations are linked. These observations may be important because although BD is characterised by episodes both of mood elevation (mania, or hypomania) and depression, it is becoming increasing recognised that there is ongoing disruption in mood and activity outside of acute mood episodes. These processes may therefore provide important mechanistic information about vulnerability to mood episodes and represent potential targets for treatment.

Mood instability has been highlighted as a risk factor for the development of BD and is associated with poorer outcomes ^1,2^. However, it is often measured retrospectively ^3^, which can introduce biases in recall due to current mood state or other factors affecting memory. Improving how we monitor, assess and define mood instability is critical for our mechanistic understanding and its potential as a treatment target in BD. The emergence of new technologies which enable the collection of digital data through smartphones, tablets, and wearable devices offers a sensitive avenue for the prospective monitoring of symptoms over time.

Circadian rest and activity pattern differences are also commonly associated with BD. Sleep problems are common ^4,5^, where insomnia or hypersomnia is associated with periods of depression and a reduced need for sleep is associated with periods of mania ^6^. These disruptions are known to persist in euthymia and periods of remission, and have been shown to encompass associated reductions in daytime activity ^7^ and greater fragmentation of daily rest-activity patterns ^8–10^. Limiting circadian rhythm disturbance may be important in bipolar disorder given evidence for a link between this disturbance and subsequent mood episode ^11,12^. Stabilising circadian rhythm disruptions has therefore been recommended in clinical guidelines13,14.

This proposed interlinked relationship has further been shown across the spectrum of mood symptoms, i.e. outside of acute mood episodes. For instance, individuals reporting periods of mood elevation and those showing depression and mania symptoms (thus, a BD phenotype) have been shown to display particularly lower objective circadian relative amplitude – a differentiation marker of daytime and night- time activity – driven by increased movement or activity during rest periods, compared to those without a BD phenotype ^5,15^. These disruptions, measured using actigraphy, remained present when individuals who went on to meet diagnostic criteria for BD were excluded ^5^, suggesting the significance of this disruption with the experience of mood dysregulation outside of particular functional impairment.

Furthermore, these patterns showed modulations with mood such that those with a history of depression had the lowest circadian amplitude and variance in this marker was more strongly associated with susceptibility to mania rather than depression ^15^.

These data suggest an interlinked relationship between mood dysregulation and disruption in circadian rest-activity patterns outside of specific diagnostic categories, but whether this co-varying relationship exists across a longitudinal scale has not been investigated. Such an investigation, emphasising temporal sensitivity in both mood instability and circadian rest-activity patterns, can aid in our mechanistic understanding of processes which may be involved in vulnerability to mood disorder as well as providing targets for future treatments.

In the current study, we tested the inter-relationship between mood and circadian-rhythm instabilities across a 10-week period in a group of individuals at low and high risk for BD (assessed by the Mood Disorder Questionnaire (MDQ) ^16^). This approach allows us to explore underlying processes involved in mood disturbance in the absence of medication confounds, which can affect both mood and circadian rhythm instability.

We used a similar approach to Rock et al., 2014, to identify individuals with experience of mood elevation, as assessed by the MDQ. We compared those with and without self-reported mood elevation on ratings of mood and symptoms at different timescales (daily vs. weekly) across a 10-week period using digital assessments made on tablets, as well as actigraphy, allowing for a finer-grained approach to identify mood instability and circadian rest-activity patterns than that provided by retrospective reports.

By focusing on the ability of remote and wearable digital technologies to provide high-quality and temporally sensitive data over a 10-week period, we hypothesised that participants scoring highly on the MDQ would show a higher level of mood instability both for positive and negative affect. We also expected the high MDQ group to show higher risk factors for BD such as higher ratings of negative affect, as well as higher levels of symptoms of mania and depression, similar to previous research ^17^. Furthermore, we predicted that those in the high MDQ group would have greater variability in their rest-activity cycle across time, as well as specific disruptions with daytime and sleep activity, as measured by actigraphy. This disruption was also predicted to be more closely associated with variability across longitudinally collected mood measures, highlighting a functional relationship between mood instability and circadian rhythm disruptions.

## Methods

### Study design

The COMET study was a prospective, between-subjects experimental cohort study over 10 weeks. All daily and weekly assessments were conducted electronically and remotely using a specialised study testing website and the True Colours online platform (www.truecolours.nhs.uk); True Colours, Department of Psychiatry, University of Oxford, Oxford, UK) on iPads (Apple Inc.) for mood assessments.

Wrist-worn GENEActiv actigraphs (ActivInsights Ltd, UK) collected actigraphy data. Data collection ran from 2015 to 2017. The study was approved by the Central University Research Ethics Committee of the University of Oxford (MSD-IDREC-C2- 2014-023).

### Participants

Participants were recruited by University of Oxford or community-wide advertisements. Seventy-four participants (48 female, 26 male; aged 18-49 years) selected based on MDQ ^16^ score gave their written informed consent to participate in the COMET study. Thirty-seven participants showed signs of mood elevation as measured by the MDQ (“high MDQ” group) (MDQ≥7) and 37 were age and gender matched, with MDQ≤5 (“low MDQ” group). In order to facilitate participant matching, those who expressed interest and met criteria for the low MDQ group were entered into a database to be invited for participation once they were matched to a high MDQ participant. We did not include the MDQ measure of functional impairment in this categorisation, similar to previous work ^17^ .

After registering interest in the study, volunteers were electronically screened using the MDQ and several questions pertaining to demographics and current or past psychiatric health. They were excluded if they did not meet criteria for the high MDQ or low MDQ groups, had a current and/or lifetime diagnosis of a DSM-IV axis I psychiatric illness (excluding BD I or II, MDD, and anxiety disorder in the high MDQ group); had a current or lifetime diagnosis of a neurological condition; and had a first degree relative with BD. Exclusion criteria also included currently using (or having done so in the past 6 weeks) lithium, antidepressant, antipsychotic, or anticonvulsant medication; or current substance abuse. The DSM-IV SCID I was used to assess all participants for current or prior psychiatric disorder at their baseline visit.

### Mood measurements

#### Daily mood assessments

Participants completed daily mood assessments 5 days per week during the 10- week study period, using the International Positive and Negative Affect Scale, Short Form (I-PANAS-SF) ^18^ . Here they were required to rate the extent to which they experienced 10 emotions on a 5-point Likert scale ranging from “very slightly/not at all” to “extremely”. Participants were asked to answer this in relation to the last 24

hours, or since they had last completed the questionnaire. The I-PANAS-SF scale measured positive affect using the words “alert”, “attentive”, “active”, “determined” and “inspired”, and negative affect using “afraid”, “ashamed”, “hostile”, “nervous” and “upset”.

#### Weekly mood assessments

Participants were also required to answer a set of three self-monitoring questionnaires as part of the online True Colours system (True Colours Team, Department of Psychiatry, University of Oxford), once a week for the 10-week study period. These questionnaires assessed mood depression using the standardised clinical scale QIDS (Quick Inventory of Depressive Symptomatology) ^20^, mood elevation using the clinical scale ASRM (Altman Self-Rating Mania scale) ^21^, and anxiety using the GAD-7 measure (Generalised Anxiety Disorder questionnaire) ^22^.

### Circadian rest-activity measurements

#### Actigraphy device

All participants were equipped with GENEActiv Original actigraphs (ActivInsights Ltd, UK), which they wore consecutively for 28 days at a time on their non-dominant wrist. To account for the 10-week study period, three watches were required to span the duration of the study. In order to facilitate battery life for each 28-day period, actigraph sampling frequency was set to 25 Hz.

#### Pre-processing of actigraphy data

Data were pre-processed using the *GGIR* package^23^ for R version 3.4.2 (R Core Team, Vienna). Non-parametric circadian-rhythm analysis (NPCRA) variables ^24^, reflecting the structure of rest-activity patterns, were calculated using the *GGIR* algorithm. Parameters included a marker for the average phase onset and activity levels for the 5-hour period of lowest activity in a given 24-hour period (L_5_), the 10- hour period of greatest activity in a given 24-hour period (M_10_), a marker for the consistency of rest-activity patterns between days (IS or interdaily stability), and a marker for within-day consolidation of rest-activity states (IV or intradaily variability). A marker of amplitude strength, measuring the difference in activity between the most

active 10-hour period (M_10_) and least active 5-hour period (L_5_), was also calculated and referred to as Relative Amplitude (RA). The equation for RA is given below.

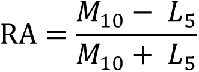

Within-subject variability of daily activity and timing (L_5_, M_10_) and RA was assessed using the timed root mean square of successive differences (*t*RMSSD) of daily measures. Further details of the actigraph device and pre-processing methods are provided in the supplementary materials.

### Statistical analysis

#### *Timed* root mean square of successive differences (*t*RMSSD)

The Root Mean Square of Successive Differences (RMSSD) ^25^ has been suggested as a measure of variability that takes into account the variance of observation-to-observation differences, as opposed to overall trial variation as measured by the standard deviation. In this sense, it takes into account gradual shifts in the mean and is particularly sensitive to temporal, or time-based, recordings. Whilst traditionally a measure used in calculating heart rate variability due to its sensitivity to high-frequency heart fluctuations ^26^, it can be particularly appropriate in the study of stability and variability in mood due to its time domain nature and sensitivity in detecting trial-by-trial fluctuations.

The equation for RMSSD is given here:

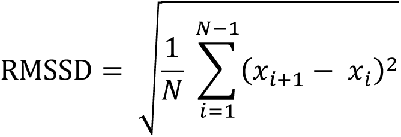

where *N* equals the number of recordings or intervals (in this case, a maximum of 51 for mood), and *x* is the value of one recording (or mood or actigraphy score). This definition, however, assumes that recordings are equally spaced in time (e.g., one recording every day at the exact same time). This is not the case with the mood data collected here wherein the five samples recorded weekly varied in time of day and consecutive intervals. Using the formula above with such variable time differences may result in artefactually high or low values of RMSSD. To address this, we use the *timed* RMSSD (*t*RMSSD) ^27^, defined as follows:

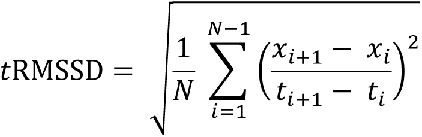

where *t_i_* corresponds to the time at which sample x_i_ was recorded. For these data, the *t*RMSSD is thus obtained by first calculating each successive difference in mood (positive affect and negative affect) over each successive time difference between recordings. This value is then squared, and the result is averaged before the square root of the total is calculated ^28^.

#### The effect of group and time on mood and circadian rest-activity patterns

General linear models (GLMs) were conducted to analyse the effect of group (high MDQ or low MDQ; the between-subjects factor), time (week; the within-subjects factor), and group-by-time interaction for I-PANAS-SF, True Colours, and actigraphy data. Separate models were conducted for weekly average and weekly variability (as measured by the *t*RMSSD) in I-PANAS-SF data, within the mood domains of positive affect and negative affect, and actigraphy data, within the rest-activity domains of L_5_, M_10_, and RA. Given that IS and IV are overall markers only and thus, daily/weekly and variability data are not available, *t*-tests were carried out to test for overall group differences in these markers. Individual GLMs were conducted for the QIDS, ASRM, and GAD-7 data, with additional post-hoc *t*-tests to test for group differences in variability (*t*RMSSD) scores. Greenhouse–Geisser corrected *p*-values are reported where appropriate, and time was *z*-scored in order to meet the assumptions of LMM analysis for actigraphy analyses. Analyses were carried out using repeated measures analyses of variance (ANOVAs) and individual sample *t*-tests in SPSS and the *lme4* package (*t*-tests using Satterthwaite’s method) (version 1.-17) ^29^ in R.

#### Relationship between daily mood monitoring, weekly mood monitoring, and circadian rest-activity patterns

Linear mixed-effects models (LMMs) were conducted to analyse the effect of group on the relationship between daily I-PANAS-SF and weekly (True Colours) mood and daily circadian rest-activity patterns across weeks, using the *lme4* package (version 1.-17) ^29^ in R. Here, mood and time were *z*-score transformed in order to meet the assumptions of LMM analysis. Both of these analyses were run on average and variability (as measured by *t*RMSSD) measures of daily mood, weekly True Colours responses, and actigraphy measures, in order for the relationship between symptomatology and mood experiences and instability to be explored.

## Results

### Participants

Seventy-four participants completed the COMET study and are included in analyses of mood data (high MDQ *n* = 37, low MDQ *n* = 37). Only those who had provided at least 50 days of actigraphy data, corresponding to the maximum of 50 days of mood data collected, are included in actigraphy analyses. Therefore, sixty-two participants are included in actigraphy analyses (high MDQ *n* = 31, low MDQ *n* = 31).

### Demographics

Overall demographic characteristics are shown in Table 1. The high MDQ group and low MDQ group were matched for age and gender. Nine participants in the high MDQ group endorsed sufficient criteria to receive a DSM-IV diagnosis, with a number of comorbidities.

**Table 1.**
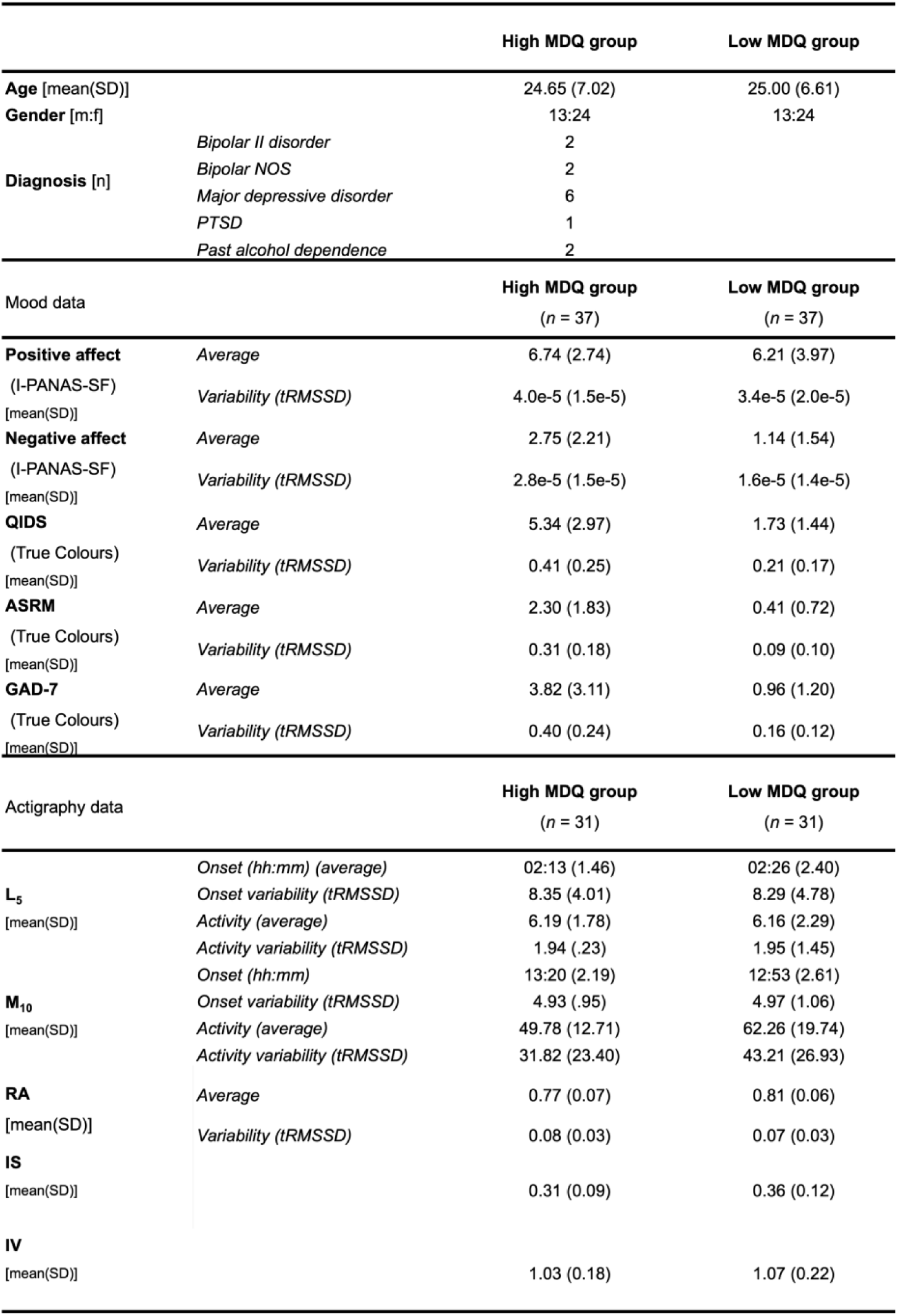
Demographics and descriptive statistics of the high MDQ and low MDQ groups.

### Number of mood recordings

There were no significant differences in the number of days of I-PANAS-SF (high MDQ: 47.65 ± 3.69, low MDQ: 49.05 ± 1.96; *U* = 550.00, *p* = .137, *r* = .17) recordings made between the high MDQ and low MDQ groups or in the number of weeks of True Colours recordings made between the two groups (high MDQ: 9.22 ± 1.47, low MDQ: 9.22 ± 1.16; *U* = 674.00, *p* = .898, *r* = .02).

### Number of actigraphy recordings

There was no significant difference in the number of days (high MDQ: 65.94 ± 6.25, low MDQ: 65.39 ± 8.57; *t*(60) = 0.29, *p* = .774) or weeks (high MDQ: 9.39 ± 0.99, low

MDQ: 9.39 ± 1.26; *U* = 467.00, *p* = .842, *r* = .00) of actigraphy data recorded between the groups.

### Daily mood assessments

Overall descriptive statistics for the daily I-PANAS-SF mood recordings are shown in Table 1.

#### Negative affect

The high MDQ group had greater average negative affect scores (*F*(1, 65) = 15.15, *p* < .001) (figure 1A), as well as greater variability in negative affect scores as measured by *t*RMSSD (*F*(1, 61) = 14.34, *p* < .001) (figure 1B). There was, however, no significant effect of time for either average or variability in negative affect (average: *F*(5.23, 339.92) = 1.72, *p* = .126; variability: *F*(5.54, 337.67) = 0.38, *p* = .882), and no group by time interaction in negative affect (average: *F*(5.54, 337.67) = 2.00, *p* = .076; variability: *F*(5.54, 337.67) = 0.85, *p* = .523).

**Figure 1.**
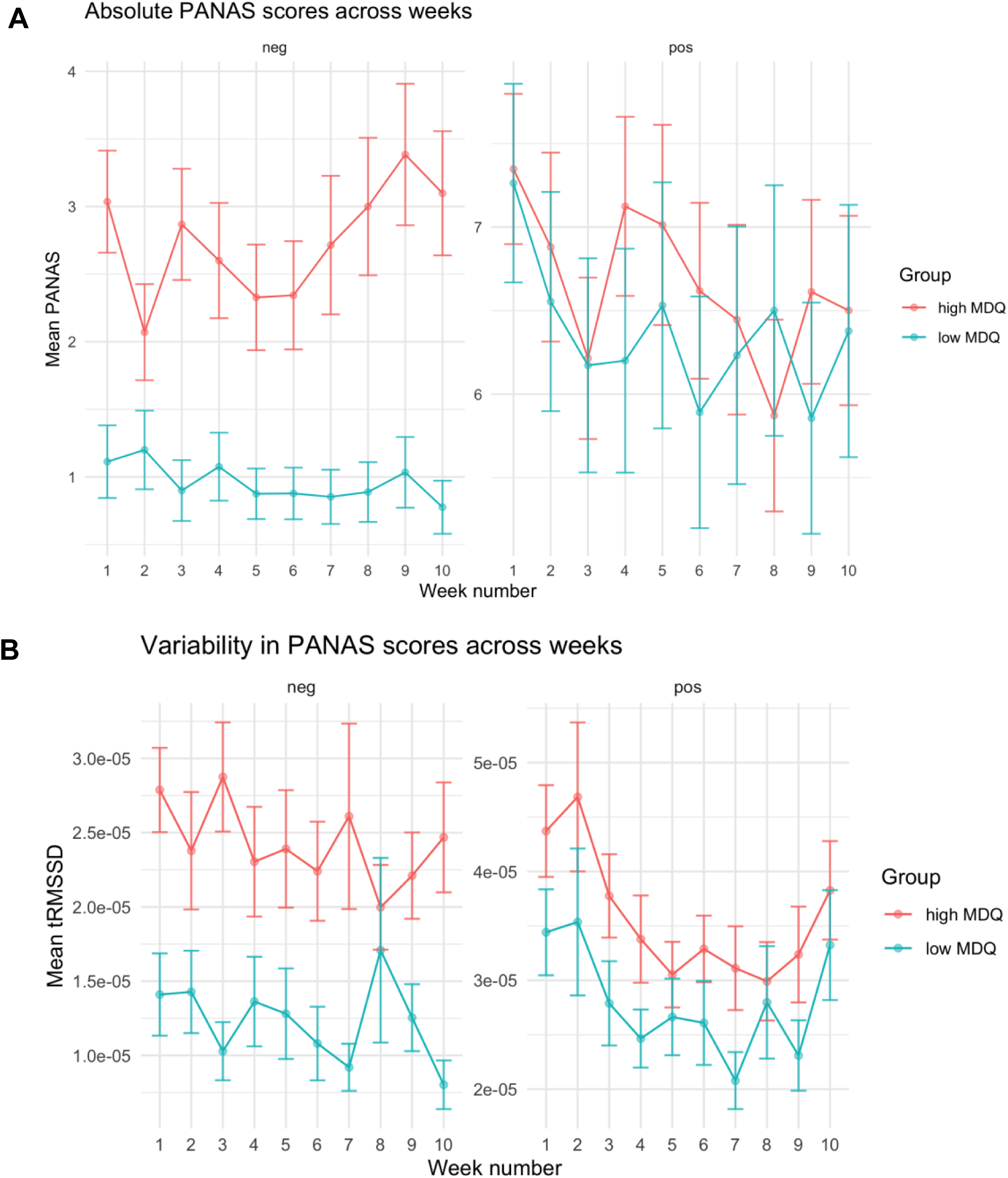
Daily I-PANAS-SF data across high MDQ and low MDQ groups. Panel A shows average negative affect across weeks (left) and average positive affect across weeks (right). Panel B shows variability in negative affect across weeks (left) and variability in positive affect across weeks (right), as measured by the *timed* Root Mean Square of Successive Differences (*t*RMSSD). Error bars represent ± 1 *SEM*.

#### Positive affect

There was no difference between groups in average positive affect scores (*F*(1, 65) = 0.05, *p* = .821) (figure 1A) but the MDQ group had greater variability in their positive affect scores compared to the low MDQ group as measured by *t*RMSSD (*F*(1, 61) = 8.18, *p* = .006) (figure 1B). An effect of time was also found, such that all participants had lower positive affect ratings across the 10-week study period (*F*(4.63, 301.21) = 2.52, *p* = .034), and also became less variable in their positive affect ratings across this period (*F*(5.15, 313.93) = 3.01, *p* = .011) but this did not interact with group (average: *F*(4.63, 301.21) = 1.04, *p* = .394; variability: *F*(5.15, 313.93) = 0.63, *p* = .679).

### Weekly mood assessments

#### Depressive symptoms

Analysis of weekly QIDS data revealed that the high MDQ group had more depressive symptoms compared to the low MDQ group (*F*(1, 41) = 21.24, *p* < .001, see Table 1) (figure 2C). Post-hoc tests on *t*RMSSD data also showed that these symptoms were more variable in the high MDQ group (0.41 ± 0.25) compared to the low MDQ group (0.21 ± 0.17) (*t*(71) = 3.98, p < .001) (figure 3). Participants also reported fewer depressive symptoms across the 10-week study period (*F*(5.69, 233.43) = 4.56, *p* < .001). An interaction analysis showed that this was different between the groups (*F*(5.69, 233.43) = 2.52, *p* = .025), such that depressive symptoms decreased in high MDQ participants to a greater degree than in low MDQ participants.

**Figure 2.**
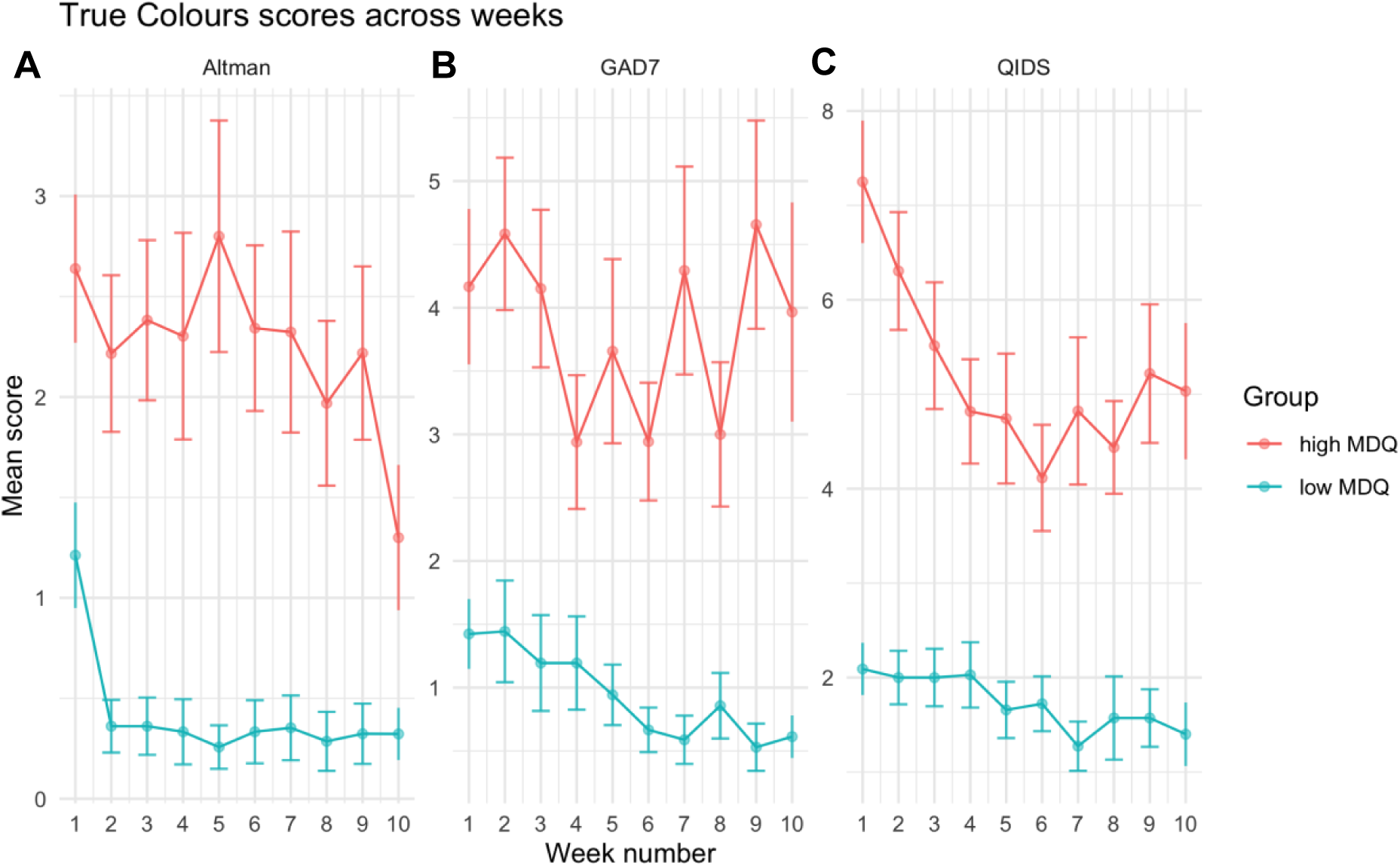
Weekly True Colours data across high MDQ and low MDQ groups. Panel A shows symptoms of mania as measured by the ASRM. Panel B shows symptoms of anxiety as measured by the GAD-7. Panel C shows symptoms of depression as measured by the QIDS. Average scores across the 10-week study period are shown. Error bars represent ± 1 *SEM*.

**Figure 3.**
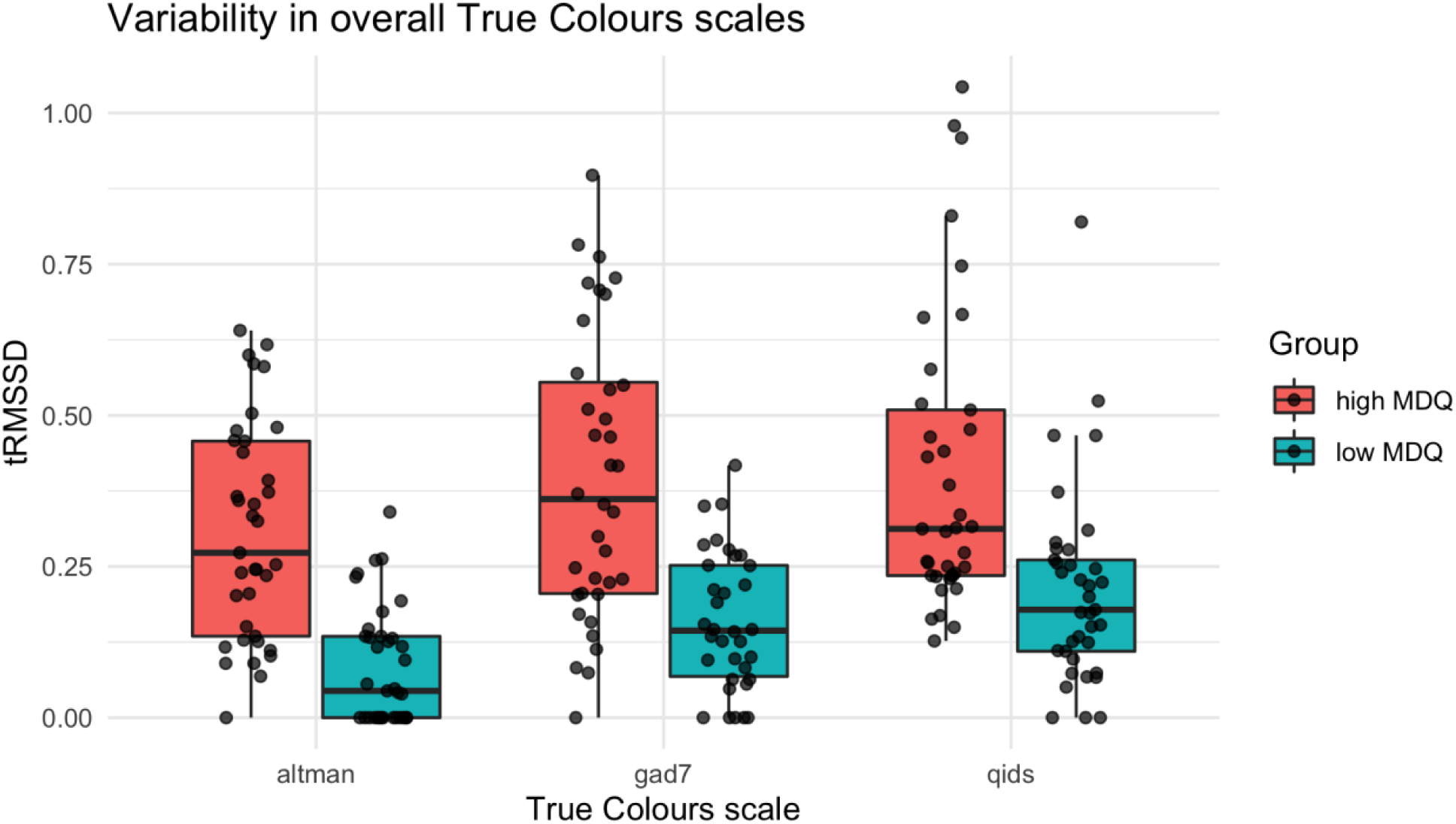
Variability in overall True Colours data as measured by the *timed* Root Mean Square of Successive Differences (*t*RMSSD), across high MDQ and low MDQ groups.

#### Manic symptoms

Analysis of weekly ASRM data revealed that the high MDQ group had more symptoms of mania compared to the low MDQ group *F(*(1, 42) = 17.11, *p* < .001, see Table 1) (figure 2A). Post-hoc tests on *t*RMSSD data also showed that these symptoms were more variable in the high MDQ group (0.31 ± 0.18) compared to the low MDQ group (0.09 ± 0.10) (*t*(71) = 6.57, p < .001) (figure 3). There was a main effect of time, such that all participants also had fewer manic symptoms over the 10- week study period (*F*(4.94, 207.66) = 3.60, *p* = .004). An interaction analysis showed that this was not different between groups (*F*(4.94, 207.66) = 1.39, *p* = .231).

#### Anxiety symptoms

Analysis of weekly GAD-7 data revealed that the high MDQ group had higher levels of anxiety compared to the low MDQ group (*F*(1, 41) = 16.25, *p* < .001, see Table 1) (figure 2B). Post-hoc tests on *t*RMSSD data also showed that these symptoms were more variable in the high MDQ group (0.40 ± 0.24) compared to the low MDQ group (0.16 ± 0.12) (*t*(67) = 3.87, p < .001) (figure 3). A main effect of time showed that anxiety tended to increase across the 10-week study period (*F*(4.01, 164.25) = 2.55, *p* = .041). An interaction analysis showed this was not different between groups (*F*(4.01, 164.25) = 2.18, *p* = .073).

### Circadian rest-activity measurements

Overall descriptive statistics for the actigraphy parameters are shown in Table 1.

Representative example actograms displaying 24-hour activity patterns in three high MDQ and three low MDQ participants are shown in figure 4. Actograms from individuals in the low MDQ group (D-F) show clear distinctions in rest-activity patterns from those of high MDQ participants (A-C).

**Figure 4.**
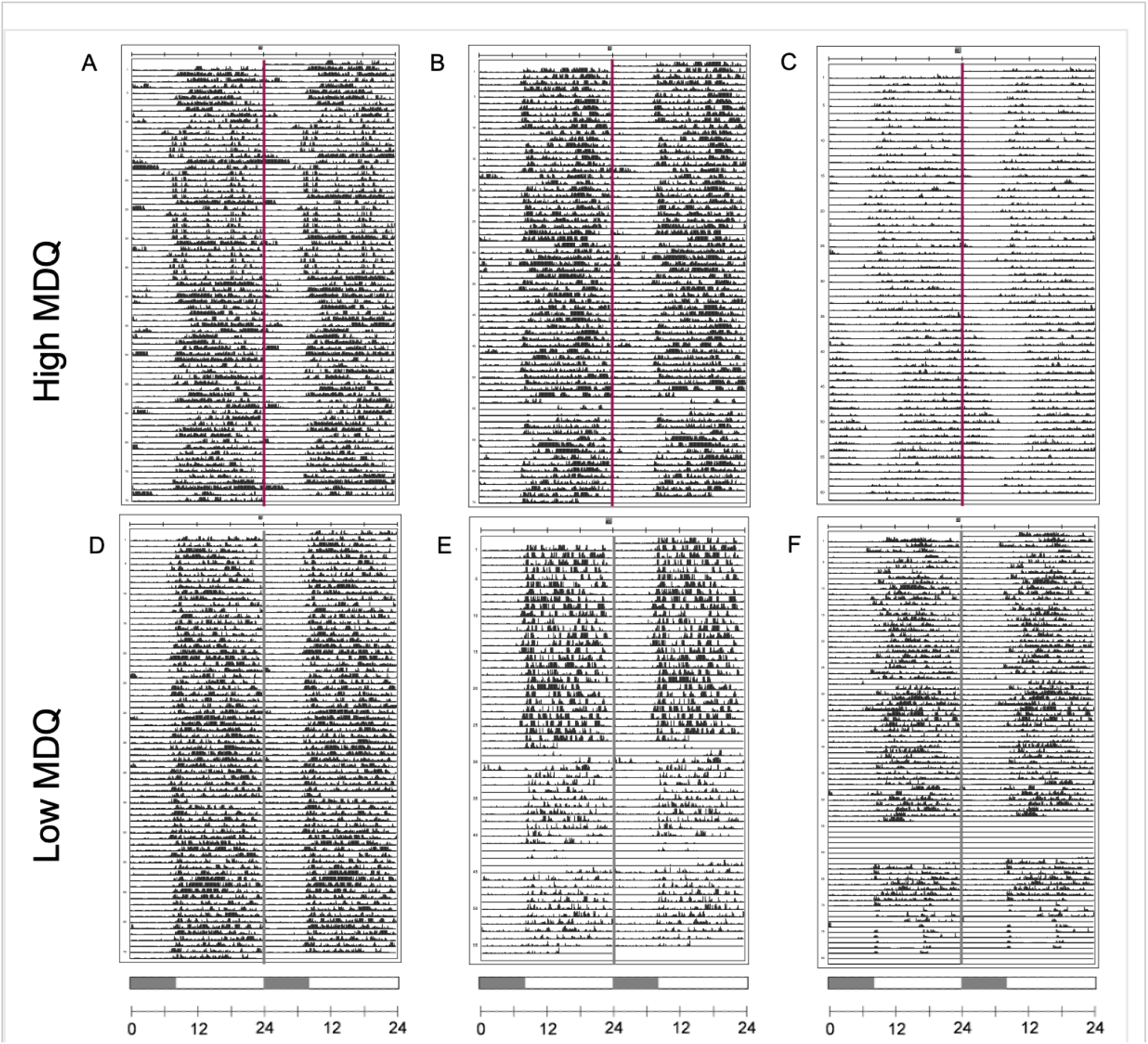
Representative actograms of rest-activity patterns. Visual inspection of double-plotted 24-h actograms generated from participants’ rest-activity patterns reflect the characteristic differences detected between groups. Shaded portion of the scale bar represents the interval between 00:00 and 08:00 h as a reference guide. Actograms were generated using ActogramJ (http://actogramj.neurofly.de/).

### L_5_

There was no difference between groups in average (*t*(62.92) = .91, *p* = .364) (figure 5A) or variability (*t*(61.75) = .25, *p* = .807) (figure 5B) in L_5_ onset. There was also no effect of time on average (*t*(502.01) = -.19, *p* = .846) or variability (*t*(494.64) = -1.64, *p* = .102) in L_5_ onset.

**Figure 5.**
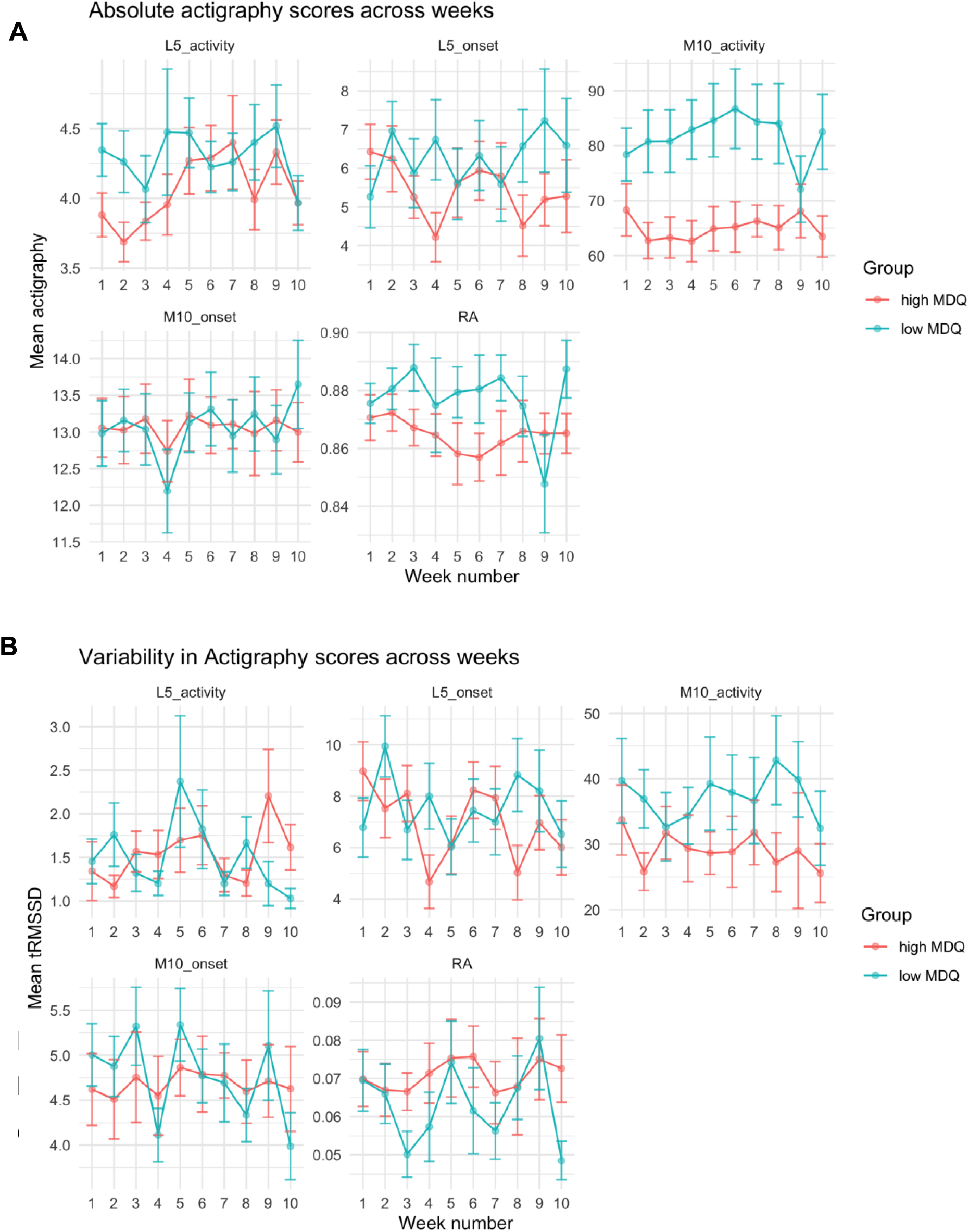
Actigraphy data across high MDQ and low MDQ groups. Panel A shows average L_5_ onset, L_5_ activity, M_10_ onset, M_10_ activity, and RA across weeks. Panel B shows variability in L_5_ onset, L_5_ activity, M_10_ onset, M_10_ activity, and RA across weeks, as measured by the *timed* Root Mean Square of Successive Differences (*t*RMSSD). Error bars represent ± 1 *SEM*.

There was no difference between groups in average (*t*(61.42) = 1.37, *p* = .177) (figure 5A) or variability (*t*(53.70) = .06, *p* = .950) (figure 5B) in L_5_ activity. While there was no effect of time on variability (*t*(496.60) = .65, *p* = .515) in L_5_ activity, there was an effect of time on average (*t*(499.77) = 2.09, *p* = .037) L_5_ activity, such that L_5_ activity tended to increase across the 10-week study period.

### M_10_

There was no difference between groups in average (*t*(61.23) = 0.05, *p* = .961) (figure 5A) or variability (*t*(61.03) = 0.46, *p* = .651) (figure 5B) in M_10_ onset. There was also no effect of time on average (*t*(498.94) = 1.18, *p* = .238) or variability (*t*(503.07) = -0.71, *p* = .479) in M_10_ onset.

The high MDQ group had lower average M_10_ activity than the low MDQ group (*t*(61.39) = 2.84, *p* = .006) (figure 5A), but there was no difference in variability in M_10_ activity between groups (*t*(59.61) = 1.52, *p* = .134) (figure 5B). There was also no effect of time on average (*t*(495.64) = 1.69, *p* = .092) or variability (*t*(48.92) = 0.01, *p* = .995) in M_10_ activity.

### RA

There was no difference between groups in average (*t*(58.83) = 1.27, *p* = .210) (figure 5A) or variability (*t*(52.41) = -1.35, *p* = .184) (figure 5B) in RA. There was also no effect of time on average (*t*(496.05) = -1.18, *p* = .239) or variability (*t*(491.87) = 0.55, *p* = .585) in RA.

### IS and IV

There was no difference between groups in average IS (*t*(60) = 1.59, *p* = .118) (figure 6A) and average IV (*t*(60) = 0.88, *p* = .383) (figure 6B).

**Figure 6.**
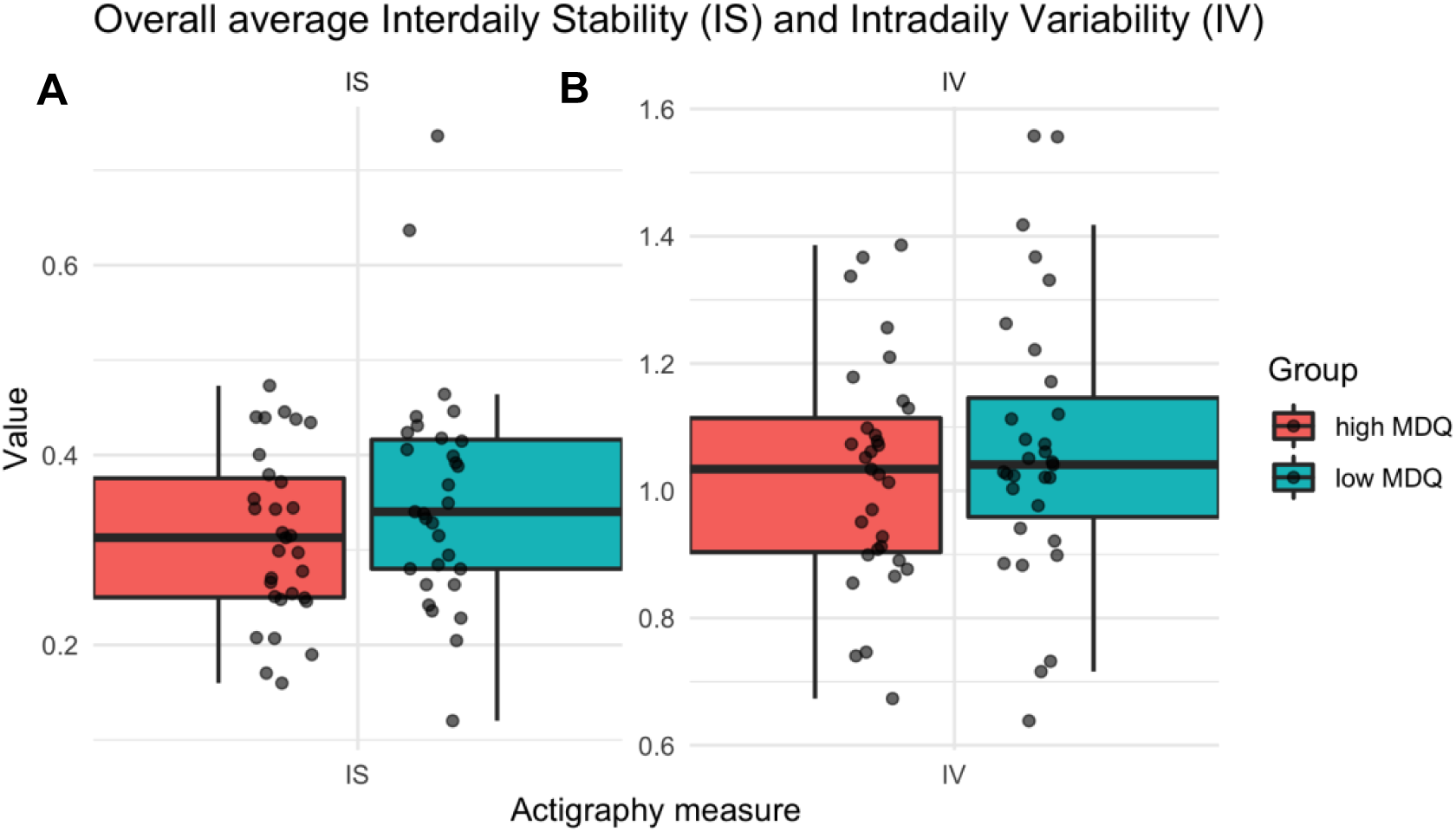
Overall interdaily stability (IS) and intradaily variability (IV) actigraphy markers, across high MDQ and low MDQ groups.

### Relationship between daily mood monitoring, weekly mood monitoring, and circadian rest-activity patterns

Exploratory linear mixed-effects modelling (LMM) was conducted to test for relationships between the actigraphy measure of interest (average M_10_ activity) and overall and variability in mood (as measured by I-PANAS-SF and True Colours) across time. There were no significant associations between mood, or mood and group interaction, on M_10_ activity in these analyses (see supplementary materials for full details).

## Discussion

The current study provides high-frequency and longitudinal information on the fluctuating nature of mood and associated patterns, supporting the use of remote and digital technologies for the development of experimental medicine models for mood instability and mood elevation. Individuals who show episodes of mood elevation as assessed by the MDQ showed differences in affect ratings and variability in ratings across different timescales. High MDQ individuals showed greater variability in their daily negative and positive mood ratings across the 10- week study period compared to matched low MDQ participants, indicating mood instability. This was further supported by increased variability in symptoms of depression, mania, and anxiety in the high MDQ group, with particular associations between mood instability and symptoms of depression and anxiety. Although the MDQ focuses on mood elevation, high MDQ individuals also reported significantly greater levels of negative affect across the 10-week study period. Together, these results support the hypothesis of increased levels of mood instability in individuals who show dimensions of BD as measured by the MDQ.

Results also support the presence of disruptions in the rest-activity cycle. The finding of lower daytime activity in the high MDQ group supports findings of disrupted circadian rest-activity patterns seen in other studies involving both patients with BD and MDD ^5,30^. While this effect also begins to suggest a wider disturbance in the mechanism responsible for regulating the cycle of daytime and night-time activity, it seems that the particular process by which this occurs varies greatly. Currently, we find that lowered daytime activity is the dominant factor in circadian disruption, whilst previous literature has reported significant disruptions in sleep indicated by greater night-time activity ^5^. Although it is true that these two mechanisms work together – that higher levels of activity during sleep or rest periods often lead to lower levels of activity during the day – the fact that we did not see this sleep-focused pattern could occur for a number of reasons.

First, the majority of previous research into this effect has included relatively short periods of monitoring (usually under 2 weeks ^3^). The current study monitored participants over 10 weeks in order to capture the dynamic nature of mood instability across time, suggesting that any sleep disruptions previously noted could be relatively acute and thus, that lowered activity during the day may be more typical of the mood disorder phenotype.

Second, along with low mood, prolonged fatigue, a lack of energy, and changes in movement and psychomotor retardation are long-established components of depression ^31^. This change in movement is particularly important to acknowledge given current findings. Considering our analysis of mood in the current sample, we know that, as well as showing mood instability, the high MDQ group were also significantly higher in negative affect and displayed greater depressive symptoms. Together, this finding of significantly lower levels of activity during the day further suggests an overall depressed mood in this group. Thus, in addition to the experience of mood instability, the combined analysis here suggests that the high MDQ group were displaying two distinct and objective symptoms of depression over the 10-week study period, both (a) absolute low mood, and (b) lower levels of movement or activity during the day.

However, the lack of an association found between concurrent mood and rest-activity patterns over the 10-week period means that we are unable to directly attribute these circadian disruptions to mood disorder symptomatology. If a relationship is indeed present, it could be that such direct consequences of fluctuating mood are symptomatic of worsening illness – and particular patterns of lowered daytime activity and disrupted sleep may be associated with clinical presentations of BD specifically ^6,7^. Future studies could account for this by focusing on specific groups of patients with mood disorders, such as those with diagnosed BD, across longitudinal periods of time in order to tease apart the nature of any dynamic relationship between mood and associated circadian rest-activity symptoms.

## Conclusion

The current study supports the use of remote and digital technologies in the diagnosis, treatment and discovery science of mood disorders, by providing high-frequency information on the fluctuating nature of mood. This is particularly important to consider given that mood instability infers a temporally sensitive, dynamic, and chronic nature. Thus, the reconceptualization of symptom management – from traditional modes of in-person clinical appointments at the temporal scale of weeks or months, to the modern use of daily and ongoing monitoring through digital technologies and passive activity sensors – is highly valuable ^32^. Current findings have increased importance in their ability to show the value of low-cost, high-compliance, digital methods of mood monitoring in the general population, with the ability to highlight distinct changes in mood and associated symptoms that may be of clinical concern.

Despite mood instability not being associated with distinct differences in circadian rest-activity rhythms, differences in these rhythms, encompassing daytime activity levels, were noted between groups. Therefore, the importance of circadian rest-activity patterns, and the utility of evolving and remote digital technologies, cannot be ignored. Circadian patterns – of sleep, daytime activity, and across the 24-hour rhythm – are increasingly considered to be core features across a range of mood disorders, including BD ^33^, and are known to be predictive of both onset and severity of illness ^34^. Given the heterogeneous nature of these patterns, the technologies used to measure them, and the mood disorder samples studied, future research should aim to assess this putative link more stringently in order to shed light on a potential homeostatic mechanism underlying mood and associated instabilities.

It is hoped that further research to identify the neural basis of this homeostatic mechanism may help explain the unique and distinct ways in which imbalance in such regulatory systems may be associated with mood disorders. Ultimately, the identification of such mechanisms will enable the development of novel treatment targets for BD and other mood disorders, optimised at regulating mood across a dynamic temporal scale.

## Supporting information

Supplementary materials including Supplemental Table 1

## Acknowledgments

This work was supported by a Wellcome Trust Strategic Award (CONBRIO: Collaborative Oxford Network for Bipolar Research to Improve Outcomes, 102,616/Z) and by the NIHR Oxford Health Biomedical Research Centre. The views expressed are those of the authors and not necessarily those of the NHS, the NIHR or the Department of Health.

