## Supplementary materials including Supplemental Table 1 for "Identifying mood instability and circadian rest-activity patterns using digital remote monitoring and actigraphy in participants at risk for bipolar disorder"

### Methods

#### Actigraphy

The actigraphs used contain a tri-axial accelerometer integrated in microelectromechanical systems and measure raw acceleration in gravitational units with a range of +/- 8g (1g = 9.8 m/s$^2$). The main output measure representing participants' moment-to-moment activity is the gravity subtracted signal vector magnitude of acceleration (Esliger et al., 2011), given by the equation:

$$\text{Acceleration magnitude} = \left(\sum \sqrt{x^2 + y^2 + z^2} - g\right)$$

Where *x*, *y*, and *z* are the three raw signals and 1g is subtracted to account for acceleration due to gravity. Negative values were rounded to zero, as per the Euclidean Norm Minus One (ENMO) (Hildebrand, Van Hees, Hansen, & Ekelund, 2014; Van Hees et al., 2014).

Only data included between the first and last midnight of actigraph recording were analysed for each participant. The *GGIR* non-wear algorithm ensured that any suspected actigraph removals were identified and missing intervals imputed with the average of similar time points from other full days of recording. Any days with more than three hours of missing data were excluded from further analyses (Bromundt et al., 2011; McGowan et al., 2019).

#### Relationship between daily mood monitoring, weekly mood monitoring, and circadian rest-activity patterns

Linear mixed-effects models (LMMs) were used to analyse the effect of group on the relationship between rest-activity patterns and mood across weeks. These analyses were conducted using the *lme4* package (version 1.-17) (Bates et al., 2015) in the R statistical programming language (R Core Team, 2022). LMMs were chosen as they allow for simultaneous estimation of between-subjects and between-recording variance, and therefore yield advantages over traditional analyses of variance (Baayen et al., 2008; Helbing et al., 2020; Kliegl et al., 2011).

The LMM was defined such that group and mood were entered as fixed effects and participant ID as the random effect, resulting in:

$$\text{rest} - \text{activity variable} \sim (mood\ \times\ group) + time +\ (1\mid participant\ ID)$$

where the interaction between group and mood, controlling for time, was tested.

The rest-activity variable of interest, average $M_{10}$ activity, was entered as the dependent variable, and the relationship with mood tested across all domains of the I-PANAS-SF (positive affect and negative affect) and True Colours (QIDS, ASRM, GAD-7) scales. Time and mood were *z*-scored in order to meet the assumptions of LMM analysis.

Within these weekly measures of rest-activity patterns and mood, both absolute (average as measured by the mean) and variability (as measured by *t*RMSSD) markers were explored, in line with the findings from the mood analyses. *P*-values were calculated using Satterthwaite's degrees of freedom method via the *lmerTest* package (version 3.1-0) (Kuznetsova et al., 2017) in R (R Core Team, 2022).

### Results

| I-PANAS-SF x $M_{10}$ activity (average) | |
|---|---|
| Negative affect (average) | $t(548.55) = 0.12$, $p = .901$ |
| Negative affect (average) x group | $t(549.08) = -0.42$, $p = .675$ |
| Negative affect (tRMSSD) | $t(513.16) = -0.86$, $p = .388$ |
| Negative affect (*t*RMSSD) x group | $t(513.08) = 0.55$, $p = .582$ |
| Positive affect *t*RMSSD | $t(504.09) = 0.07$, $p = .947$ |
| Positive affect (*t*RMSSD) x group | $t(504.08) = -0.06$, $p = .951$ |
| **True Colours x $M_{10}$ activity (average)** | |
| QIDS (average) | $t(501.57) = -0.50$, $p = .615$ |
| QIDS (average) x group | $t(502.51) = 0.56$, $p = .579$ |
| ASRM (average) | $t(491.42) = 0.40$, $p = .693$ |
| ASRM (average) x group | $t(493.16) = 0.13$, $p = .900$ |
| GAD-7 (average) | $t(496.71) = 0.22$, $p = .825$ |
| GAD-7 (average) x group | $t(494.79) = -0.58$, $p = .562$ |

Supplementary Table 1. Results of the linear mixed-effects models (LMMs), testing the association between mood on average $M_{10}$ activity over time. Mood by group interaction effects are also shown.
